# Quantifying absolute gene expression profiles reveals distinct regulation of central carbon metabolism genes in yeast

**DOI:** 10.1101/2020.12.22.423946

**Authors:** Rosemary Yu, Egor Vorontsov, Carina Sihlbom, Jens Nielsen

**Affiliations:** Department of Biology and Biological Engineering, Chalmers University of Technology, SE-412 96 Gothenburg, Sweden; Novo Nordisk Foundation Center for Biosustainability, Chalmers University of Technology, SE-412 96 Gothenburg, Sweden; Proteomics Core Facility, Sahlgrenska Academy, University of Gothenburg, SE-413 90 Gothenburg, Sweden; Novo Nordisk Foundation Center for Biosustainability, Technical University of Denmark, DK-2800 Kgs. Lyngby, Denmark; BioInnovation Institute, Ole Måløes Vej 3, DK-2200 Copenhagen N, Denmark

## Abstract

In addition to specific regulatory circuits, gene expression is also regulated by global physiological cues such as the cell growth rate and metabolic parameters. Here we examine these global control mechanisms by analyzing an orthogonal multi-omics dataset consisting of absolute-quantitative abundances of the transcriptome, proteome, and intracellular amino acids in 22 steady-state yeast cultures. Our model indicates that transcript and protein abundance are coordinately controlled by the cell growth rate via RNA polymerase II and ribosome abundance, but are independently controlled by metabolic parameters relating to amino acid and nucleotide availability. Genes in central carbon metabolism, however, are regulated independently of these global physiological cues. Our findings can be used to augment gene expression profiling analyses in the distantly related yeast *Schizosaccharomyces pombe* and a human cancer cell model. Our results provide a framework to analyze gene expression profiles to gain novel biological insights, a key goal of systems biology.

## Introduction

In the presence of genetic or environmental perturbations, differential expression of genes, orchestrated by dedicated regulatory circuits, shapes the physiological responses of the cell. Common physiological responses to perturbations, e.g. in response to stress or during oncogenic transformation, often include changes in the cell growth rate and metabolism. In turn, both growth rate and metabolic parameters of the cell can exert global influences on gene expression, as demonstrated by landmark studies in *E. coli* (*1–4*) and in yeast (*5–8*). Thus, the gene expression program following a perturbation reflects a joint effect of the specific regulatory circuits that are induced (or repressed) by the perturbation, as well as the global influence on gene expression by an altered physiological state (Fig 1A). Further complexities arise as gene expression and cell physiology operate in mutual feedback (*4, 9*), which can lead to the emergence of complex behaviours (*10*). Currently, a quantitative framework to understand the global effects of cell physiology on gene expression is lacking. Development of such a framework would allow perturbation-specific gene regulation mechanisms to be uncoupled from global gene expression control, and allow synthetic gene circuits with complex behaviours to be designed (*9, 10*).

**Fig 1.**
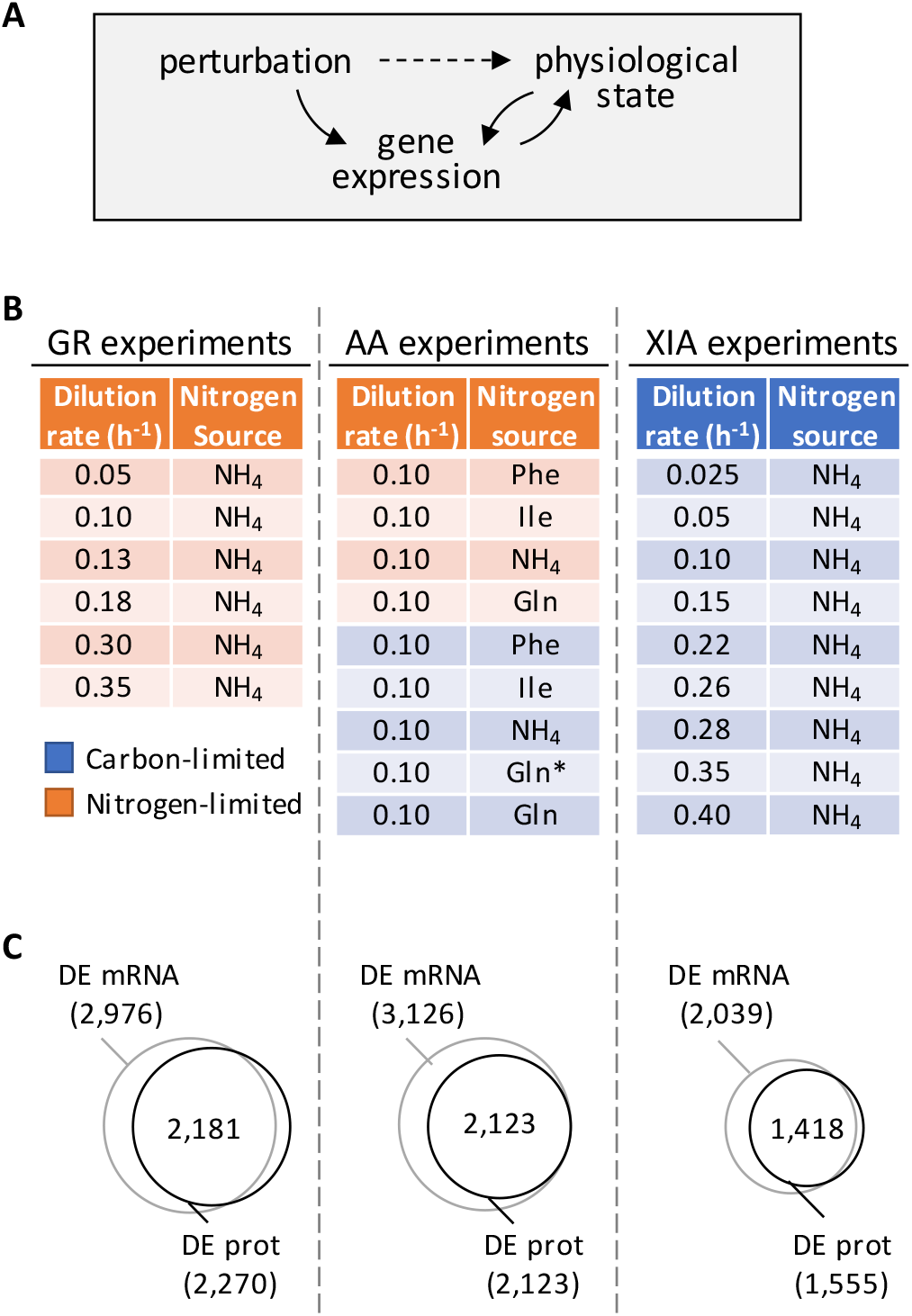
Global regulation of gene expression by the physiological state of the cell. A. Gene expression profiles of a cell depends on both specific gene expression programs induced by specific perturbations, as well as the global influence on gene expression by the physiological state of the cell. B. Experimental design to orthogonally probe the effects of growth rate and amino acid metabolism on gene expression. Cells were grown in chemostats at controlled growth rates and media composition. GR, growth rate; AA, amino acid; XIA, Xia *et al* (*22,23*). In AA experiments, carbon-limited conditions (blue rows), the “Gln” condition and the “Gln*” condition differ in the concentration of Gln and glucose in the chemostat feed media; see Table S1 for full details. C. Number of differentially expressed (DE; FDR < 0.01) genes at mRNA and protein (prot) levels in the GR experiments, AA experiments, and XIA experiments, showing that a large number of genes are regulated by growth rate and metabolic parameter.

Seminal studies in the field (*11–13*) have previously examined the interaction between growth rate, metabolic parameters, and gene expression, using microarrays and relative-quantitative metabolomics, in the eukaryal model organism *Saccharomyces cerevisiae.* Herein we revisit these interactions using RNAseq-based absolute-quantitative transcriptomics, showing substantial changes in absolute quantities of mRNA between different growth conditions which cannot be captured with relative-quantitative data. We further provide absolute-quantitative proteomics and intracellular amino acid abundance, in a total of 22 steady-state yeast cultures in biological triplicates, as a high-quality resource to the community. The 22 steady-state conditions were designed to orthogonally probe the effects of growth rate and metabolic parameters related to amino acids on gene expression (Fig 1B). We found that ~90% of genes are globally influenced by the cell growth rate and/or metabolic parameters. The growth rate-induced gene expression changes were coordinated at the transcript and protein levels, and were associated with the availabilities of the transcription and translation machineries. In contrast, gene expression control by metabolic parameters were not associated with the availability of transcription and translation machineries, but were likely regulated by the availabilities of amino acids and nucleotides. We found that genes related to central carbon metabolism (CCM) were distinctly regulated, reflecting unique control mechanisms to ensure robust expression of this metabolic pathway. Finally, by re-analyzing gene expression profiles of a distantly related yeast, *Schizosaccharomyces pombe,* and of the human Burkitt’s lymphoma cell line P493-6, we demonstrated that our findings can be broadly applied to uncouple global gene expression control from regulation by specific transcriptional and translational circuits, allowing novel biological insights in gene regulation to be uncovered.

## Results

### Data description

We performed a series of 14 steady-state chemostat cultures of *S. cerevisiae* (Fig 1B and Table S1) which orthogonally modulated either the growth rate (GR experiments) under nitrogen limitation, or the amino acid metabolic parameters (AA experiments) under either nitrogen or carbon limitation. The nitrogen sources in the AA experiments were chosen to represent those that are preferred (NH4 and Glu) and non-preferred (Phe and Ile), as previously defined (*14*). Depending on the growth condition, the total protein content in the dry cell weight ranged between 20.6% and 59.4%, and the total RNA content ranged between 1.8% and 8.9%, (Table S1), in agreement with previous observations of total protein and RNA content with changing growth rate and nitrogen source (*8,15, 16*). We then profiled the absolute-quantitative transcriptomic and proteomic abundances (mmol/gDW) of these chemostat cultures in biological triplicates, generating a multi-omics dataset containing 3,127 transcript-protein pairs across 14 conditions (Table S2). In both the GR experiments and the AA experiments, we found >2,000 genes (68-70%) to be differentially expressed at both the transcript and protein levels with FDR < 0.01 (Fig 1C), indicating that gene expression is subject to extensive global control by growth rate and metabolic parameters. A total of 1,493 genes (48%) in the GR experiments, and 1,959 genes (63%) in the AA experiments, were differentially expressed not only significantly, but by more than two-fold. In Supplemental Fig 1 we present the total protein-RNA correlation of these chemostat cultures, with a Pearson *r* of 0.51 and a Spearman *ρ* of 0.27, and sample-wise proteome-transcriptome correlations, with a median Pearson *r* of 0.40 and Spearman *p* of 0.63; agreeing well with previous studies (*8, 17–19*). In Supplemental Fig 2, we present the measured absolute quantities of subunits for several complexes (*8, 20*), showing that our measured protein abundances agree with the known subunit stoichiometry within 1-2 orders of magnitude, typical of iBAQ-based proteomics quantitation (*8, 18, 21*). This variability in the proteomics data reflects a mixed effect of inaccuracies in the proteomics methodology, and proteins that are partially synthesized, partially degraded, and not being part of its dessignated protein complex. For f1f0 ATP synthase, ATP15 and TIM11 were 4 orders of magnitude lower than the abundance of other subunits (Supplemental Fig 2), likely reflecting the difficulty in extracting and quantifying membrane-embedded proteins.

We further mined absolute-quantitative transcriptomics and proteomics data from Xia *et al (22, 23*) (XIA experiments), wherein *S. cerevisiae* were grown at 9 different dilution rates with glucose being the limiting nutrient (Fig 1B-C). Data from the XIA experiments, containing paired transcript-protein abundance of 2,235 genes (Table S3), allowed us to validate our findings in the GR experiments in a different nutrient limitation setting.

### Global regulation of transcription by growth rate and metabolic parameters

We first considered the global control of mRNA abundance by growth rate and metabolic parameters. We used 25 clustering indices (*24*) to determine the best clustering scheme for transcript abundance in the GR experiments, and found that they fell into an optimal number of 3 clusters (Fig 2A). The list of genes in each of the clusters can be found in Table S2 and filtering by the column “GR.cluster”. Expression of transcripts in GR.cluster.1 and GR.cluster.3, together including 2,895 genes (92%), exhibited strong associations with growth rate (Fig 2B). In contrast, expression of 232 transcripts (7%) in GR.cluster.2 did not increase with increasing growth rate (Fig 2C). These growth rate-independent genes in GR.cluster.2 were enriched in GO-slim terms (*25*) related to CCM, including “carbohydrate metabolic process” and “generation of precursor metabolites and energy” (Fig 2D). We further observed that genes in GR.cluster.1 and GR.cluster.3 approximately followed the protein abundance of RNA polymerase II (*26*) (Fig 2E), suggesting that transcript abundance of these 92% of genes largely reflected the availability of RNA polymerase II, consistent with previous studies (*4, 27, 28*). In contrast, the 232 genes in GR.cluster.2 exhibited slightly decreased abundance with increasing RNA polymerase II expression (Fig 2E), indicating that these genes were regulated by a mechanism independent from growth rate-associated changes in RNA polymerase II.

**Fig 2.**
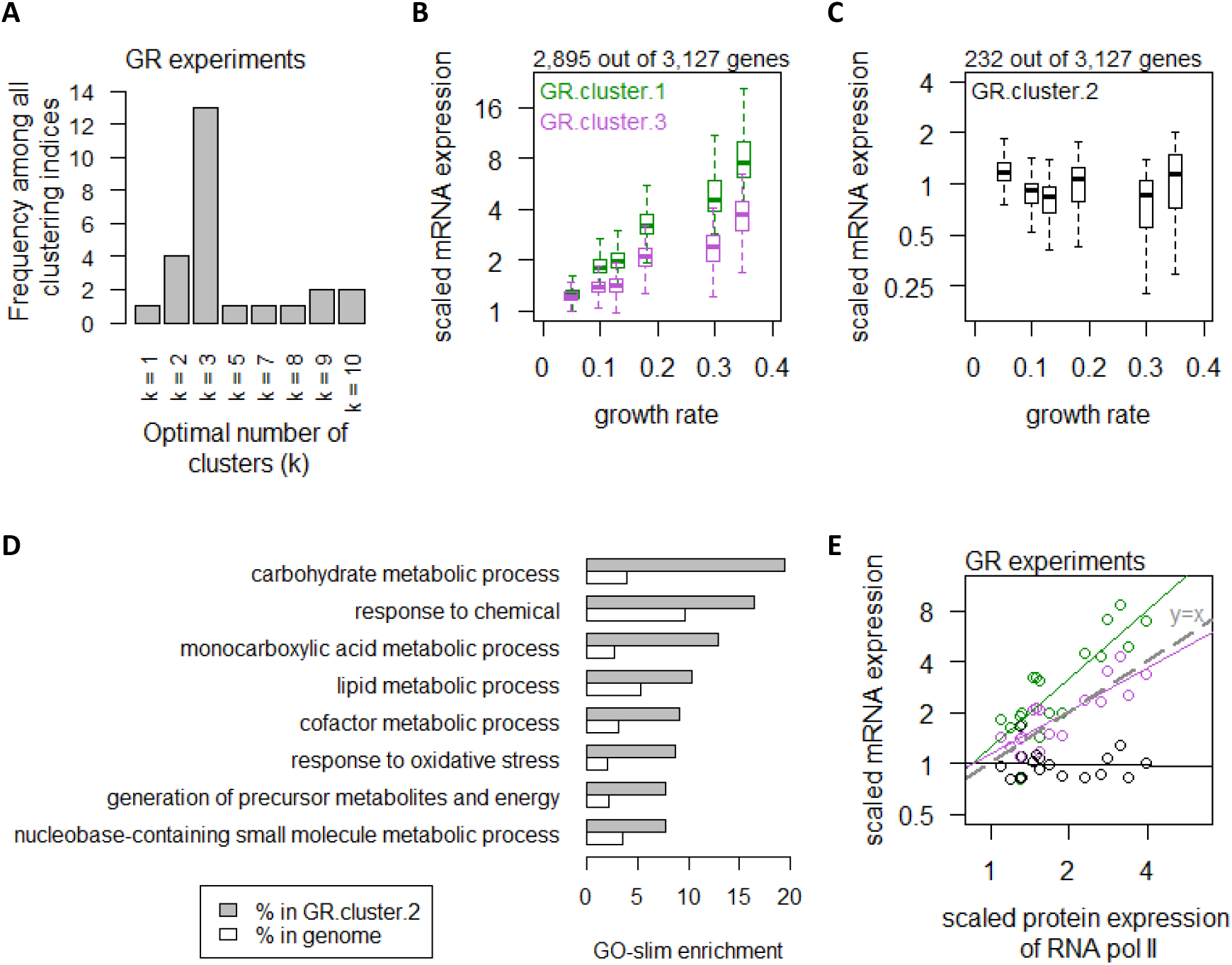
Global control of transcript abundance by growth rate. A. Using 25 clustering indices, we found that most indices suggest an optimal number of 3 clusters for transcript abundance in the GR experiments. B. Abundance of transcripts in GR.cluster.1 (green) and GR.cluster.3 (purple). Center line, median; box limits, upper and lower quartiles; whiskers, 1.5x interquartile range. C. Abundance of transcripts in GR.cluster.2. Center line, median; box limits, upper and lower quartiles; whiskers, 1.5x interquartile range. D. GO-slim enrichment of genes in GR.cluster.2 showing enrichment in GO-slim terms related to CCM, among others. E. Expression of RNA polymerase II protein abundance and mRNA abundance of the three clusters in the GR experiments. Colors are as panels B-C. Median mRNA expression values in each cluster and median protein expression of RNA polymerase II are shown. Grey dashed line represents y=x.

To confirm these findings, we performed similar analyses using data from the XIA experiments (*22, 23*), where the optimal number of clusters was calculated to be 2 clusters (Supplemental Fig 3A). The list of genes in each of the clusters in the XIA experiment can be found in Table S3 and filtering by the column “XIA.cluster”. XIA.cluster.2 contained 1,964 genes (88%) which exhibited growth rate-dependent transcript abundance (Supplemental Fig 3B), while XIA.cluster.1 contained 271 genes (12%) showing growth rate-independence. Genes in XIA.cluster.1 were enriched in CCM GO-slim terms (Supplemental Fig 3C-D), including “carbohydrate metabolic process” and “generation of precursor metabolites and energy”, validating our results in the GR experiments. Of note, GO-slim terms related to CCM (“carbohydrate metabolic process”; “cellular respiration”; and “generation of precursor metabolites and energy”) were enriched among the most abundant 10% genes in both datasets (Supplemental Fig 3E-F), demonstrating that the transcript expression of vast majority of genes were controlled by the cell growth rate, but a distinct mechanism regulated a small number of CCM-related transcripts which are highly abundant.

In the AA experiments, using the same 25 indices (*24*) we calculated that transcript expression fell into an optimal number of 2 clusters (Fig 3A). The list of genes in each of the clusters in the AA experiments can be found in Table S2 and filtering by the column “AA.cluster”. In AA.cluster.1, a total of 3,011 genes (96%) exhibited increasing expression levels when cells were grown on preferred nitrogen sources (NH4 and Gln) compared to non-preferred nitrogen sources (Phe and Ile), under carbon-limiting conditions (Fig 3B). In contrast, 116 transcripts (4%) in AA.cluster.2 exhibited consistent expression levels regardless of the amino acid metabolic parameters (Fig 3C). These genes were enriched in processes related CCM, as well as amino acid metabolism (Fig 3D). Taken together, these results indicate that regulation of CCM-related genes was distinct from global gene expression control by both growth rate and metabolic parameters of the cell.

**Fig 3.**
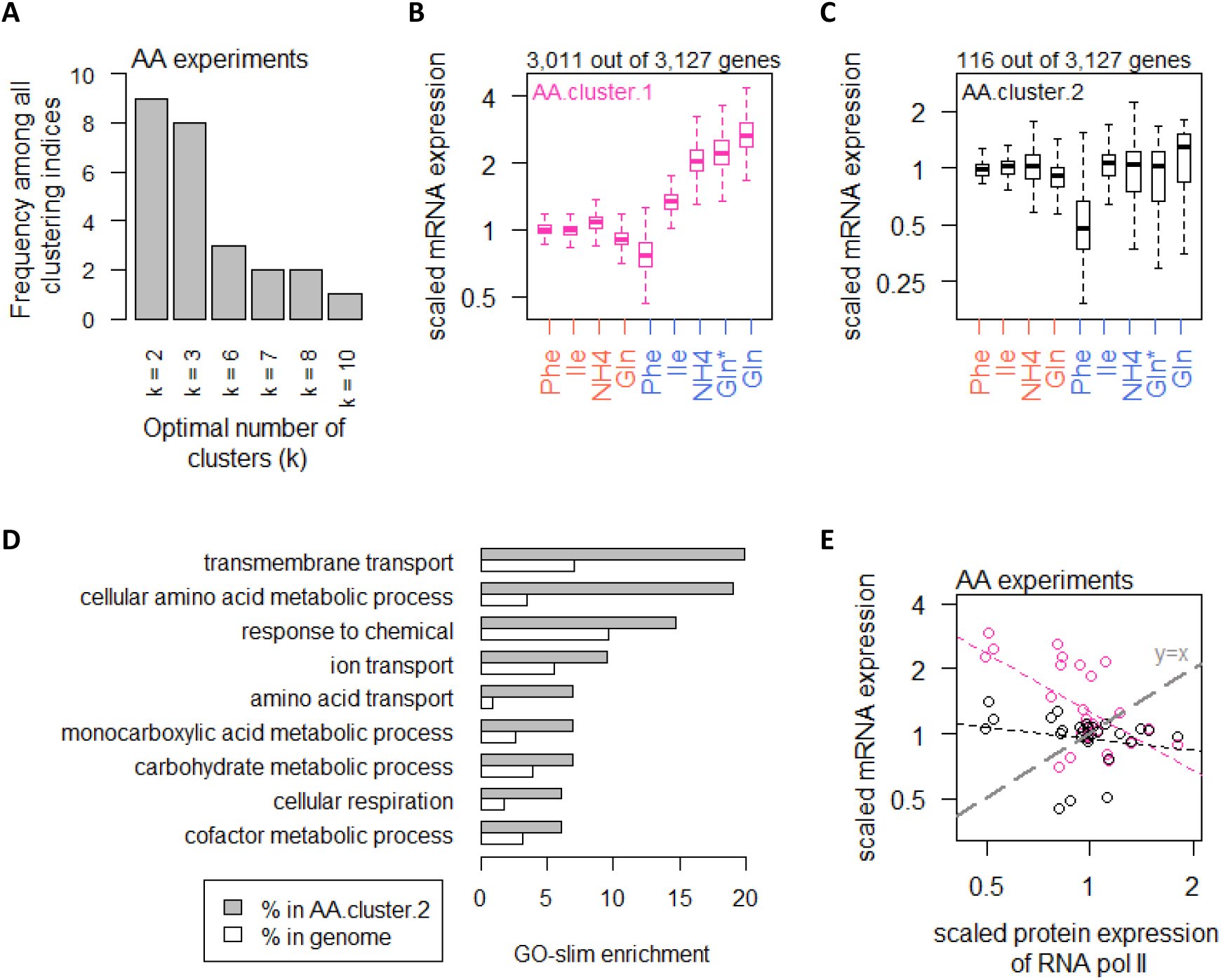
Global control of transcript abundance by amino acid metabolic parameters. A. Using 25 clustering indices, we found that most indices suggest an optimal number of 2 clusters for transcript abundance in the AA experiments. B. Abundance of transcripts in AA.cluster.1. Center line, median; box limits, upper and lower quartiles; whiskers, 1.5x interquartile range. In AA experiments, carbon-limited conditions (blue rows), the “Gln” condition and the “Gln*” condition differ in the concentration of Gln and glucose in the chemostat feed media; see Table S1 for full details. C. Abundance of transcripts in AA.cluster.2. Center line, median; box limits, upper and lower quartiles; whiskers, 1.5x interquartile range. In AA experiments, carbon-limited conditions (blue rows), the “Gln” condition and the “Gln*” condition differ in the concentration of Gln and glucose in the chemostat feed media; see Table S1 for full details. D. GO-slim enrichment of genes in AA.cluster.2 showing enrichment in GO-slim terms related to CCM, among others. E. Expression of RNA polymerase II protein abundance and mRNA abundance of the two clusters in the AA experiments. Colors are as panels B-C. Median mRNA expression values in each cluster and median protein expression of RNA polymerase II are shown. Grey dashed line represents y=x.

Examining RNA polymerase II levels in the AA experiments, we found that changes in transcript abundance did not reflect RNA polymerase II availability in response to metabolic parameters (Fig 3E). As the increased transcripts in the majority of genes when cells were grown on preferred nitrogen sources (Fig 3B) must be supported by increased biosynthesis of nucleotides, we therefore examined the intracellular concentrations of Ser and Gly, which are substrates for purine synthesis (Fig 4A), as well as Gln and Asp, which are upstream of both purine (Fig 4A) and pyrimidine synthesis (Fig 4B). We found that transcript abundance in the AA experiments tracked closely with intracellular Ser and Gly concentrations (Fig 4C), but not with Gln or Asp (Fig 4D). Moreover, in absolute-quantitative terms (Table S4), Gln and Asp were present at much higher intracellular concentrations (mean of 166.7 and 12.4 μmol/gDW, respectively, across all AA experiments) compared to Gly and Ser (mean of 1.5 and 2.8 μmol/gDW, respectively, across all AA experiments). Together, this suggests that intracellular Gly and Ser were likely limiting for nucleotide synthesis, particularly for purines, which thereby regulated global transcript abundance when growth conditions change.

**Fig 4.**
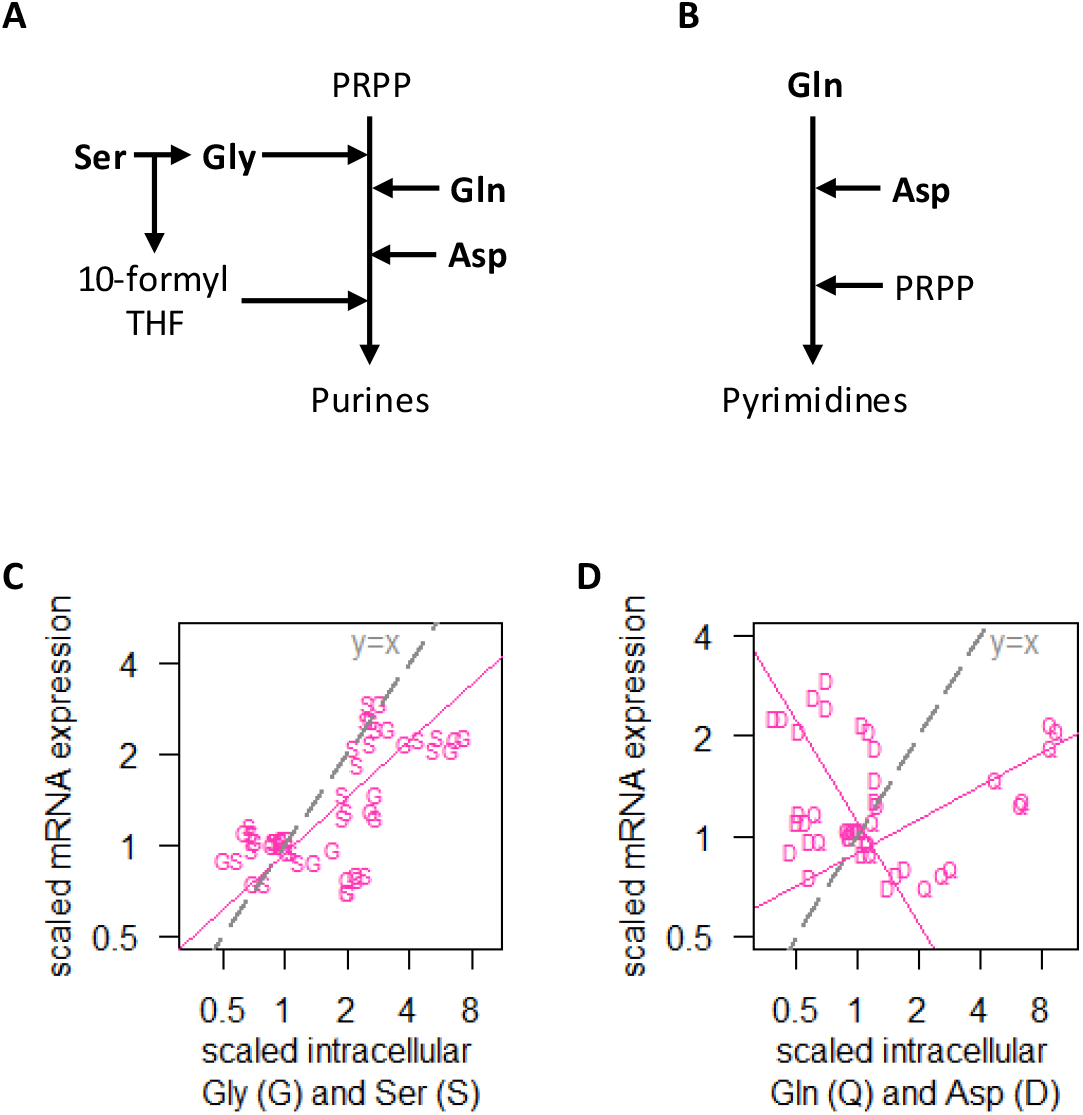
Global control of transcript abundance by amino acid metabolic parameters. A. Simplified pathway of purine synthesis. B. Simplified pathway of pyrimidine synthesis. C. Intracellular concentrations of Gly (G) and Ser (S), and RNA abundance of genes in AA.cluster.1, showing that transcript abundance for most genes in the AA experiments track closely with intracellular Ser and Gly concentrations. Median mRNA expression values of AA.cluster.1 are shown. D. Intracellular concentrations of Gln (Q) and Asp (D), and RNA abundance of genes in AA.cluster.1, showing that transcript abundance for most genes in the AA experiments do not track with intracellular Gln and Asp concentrations. Median mRNA expression values of AA.cluster.1 are shown.

### Global regulation of translation by growth rate and metabolic parameters

We next examined the correlation between mRNA and protein abundance of each gene, either in the GR experiments alone; in the AA experiments alone; or with data from the two experiments combined. As the underlying distribution of the mRNA and protein abundances in these analyses were non-normal (*p_Shapiro-Wilk_* < 0.01) for a large portion of genes, we used Spearman correlations in the following analyses. We note that Spearman correlation coefficients (*ρ*) were closely related to Pearson correlation coefficients (*r*) in nearly all of our analyses (Supplemental Fig 4), and as such our observations can be easily compared to previous studies which have used Pearson correlations. In Table S2 we report both Spearman *ρ* and Pearson *r* values.

Several previous studies have suggested that, comparing the mRNA and protein abundances of the same genes across different experimental conditions, generally there is a high correlation between protein and mRNA abundance (*17, 29–31*). Here, in the GR experiments, we observed high protein-mRNA correlations consistent with these previous studies, with a median *ρ* of 0.76 (Fig 5A) and 2,379 genes (78%) having *ρ* > 0.5. Calculating the protein translation rate for each gene (*k_sP_*, in the unit of protein/mRNA/h; see Methods section for full details) (*17, 18*), we found that overall *k_sP_* is increased with increasing growth rate (Fig 5B). Concurrently we observed an increase in the % (mol/mol) of ribosomal proteins in the total proteome (Fig 5C), consistent with previous reports (*29, 32*) and suggesting that the increase in *k_sP_* in the GR experiments were linked to the availability of ribosomes. Taken together, our data indicate that increasing growth rate modulated global gene expression via coordinated increases in both transcription and translation, by increasing the availability of RNA polymerase II and ribosomes.

**Fig 5.**
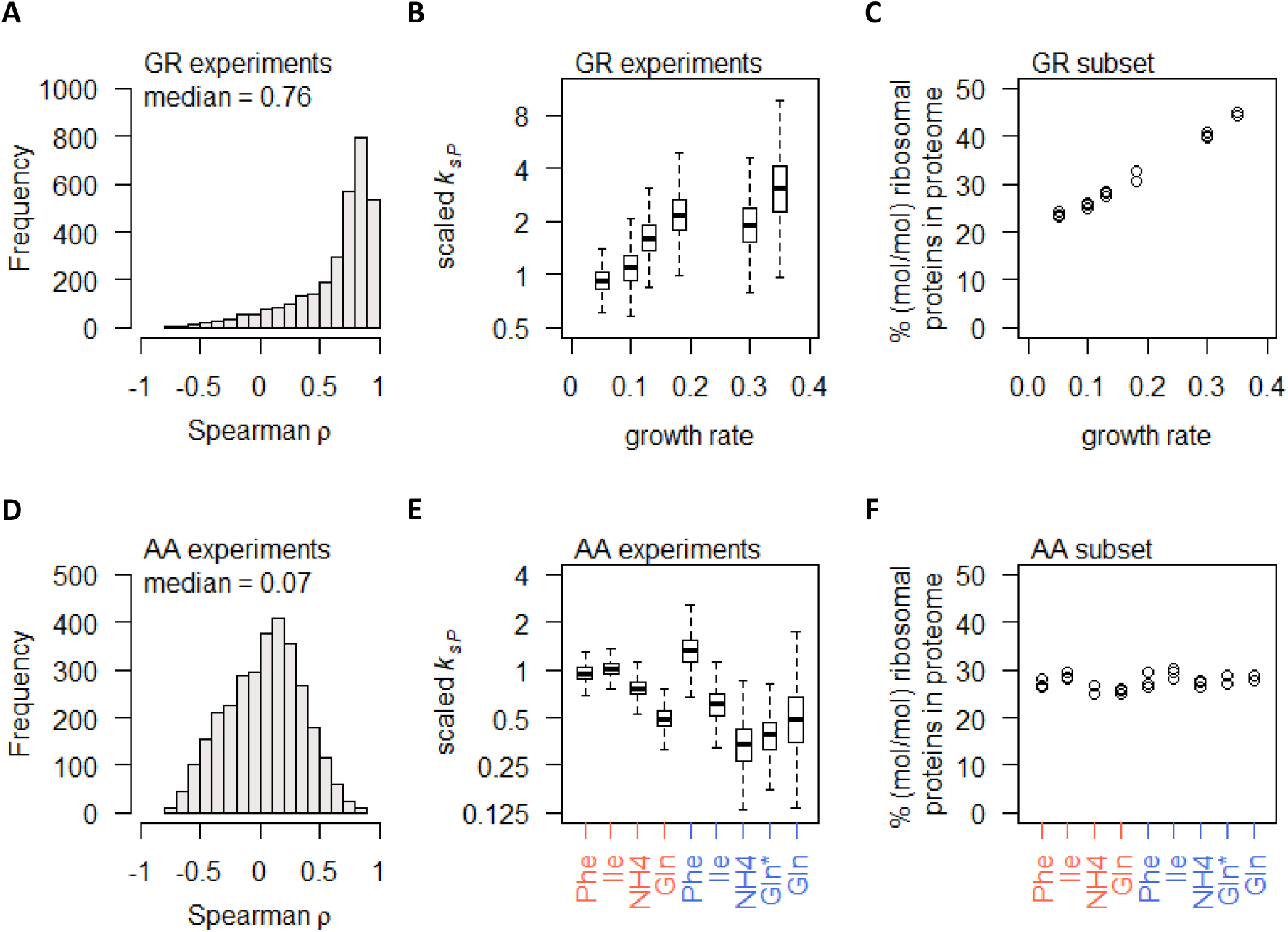
Global control of protein synthesis by growth rate and amino acid metabolism. A. Spearman correlation of absolute mRNA and protein abundances for each gene is calculated in the GR experiments, and the distribution is shown demonstrating overall high correlation. B. Protein translation rate (*k_sP_*) in the GR experiments increases with growth rate by about 4-fold overall. Center line, median; box limits, upper and lower quartiles; whiskers, 1.5x interquartile range. C. Relative ribosomal protein abundance in the GR experiments was calculated as the sum of all detected ribosomal proteins and normalized to the total protein content, showing a linear increase with increasing growth rate. D. Spearman correlation of absolute mRNA and protein abundances for each gene is calculated in the AA experiments, and the distribution is shown demonstrating overall poor correlation. E. Protein translation rate (*k_sP_*) in the AA experiments decreases when cells were grown on preferred nitrogen sources (NH4 and Gln) compared to non-preferred nitrogen sources (Phe and Ile). In N-limited cultures (orange) there is an overall 2-fold decrease, and in C-limited cultures (blue) there is an overall 4-fold decrease. Center line, median; box limits, upper and lower quartiles; whiskers, 1.5x interquartile range. In C-limited conditions (blue), the “Gln” condition and the “Gln*” condition differ in the concentration of Gln and glucose in the chemostat feed media; see Table S1 for full details. F. Relative ribosomal protein abundance (sum of all detected ribosomal proteins normalized to total protein) is constant in the AA experiments, where the growth rate is controlled to a constant as seen in Fig 1B. In C-limited conditions (blue), the “Gln” condition and the “Gln*” condition differ in the concentration of Gln and glucose in the chemostat feed media; see Table S1 for full details.

In contrast to the GR experiments, the gene-specific protein-mRNA correlations in the AA experiments were surprisingly poor, with a median *ρ* of 0.07 (Fig 5D) and most genes with −0.5 < *ρ* < 0.5 (2,763 genes, 88%). To check whether the poor correlations stem from lower range of differential gene expression, we split the total of 3,127 genes into quartiles by mRNA differential expression (fold-change of largest to smallest measured value) and found that, even in the 4^th^ quartile where the range of differential expression was >5.3-fold, the protein-mRNA correlations remained very low, with a median *ρ* of 0.24 (Supplemental Fig 5A). Similar results were found when genes were split into quartiles by protein differential expression (Supplemental Fig 5B), indicating that the poor protein-mRNA correlations observed for most genes in the AA experiment cannot be explained by the range of differential gene expression alone. Calculating the protein translation rate *k_sP_*, we found that gene-specific *k_sP_* in the AA experiments were globally reduced when cells were grown on preferred nitrogen sources, in both nitrogen-limited cultures and in carbon-limited cultures (Fig 5E). These global changes in *k_sP_* occurred without any changes in the % of ribosomal proteins in the total proteome (Fig 5F). As the growth rate of the AA experiments were controlled to a constant 0.1 h^-1^ (Fig 1B), this indicates that the % of ribosomal proteins in the total proteome was a direct reflection of the cell growth rate, while changing metabolic parameters modulated protein translation rates without modifying the abundance of ribosomes.

Considering all gene expression measurements in GR and AA experiments combined, the protein-mRNA correlations remained poor, with a median *ρ* of 0.22 (Fig 6A). To further examine these protein-mRNA relationships, we tested for enrichment of GO-slim terms in 200-gene sliding windows of increasing *ρ* (*8, 19*). We found that genes with high correlations between protein and mRNA abundance (median *ρ* of 0.4 to 0.7 in 200-gene windows) are enriched in GO-slim terms related to amino acid metabolism, as well as processes related protein translation, including ribosome biogenesis; rRNA processing; tRNA aminoacylation; and so on (Fig 6B). This indicates that processes related to protein translation were themselves regulated transcriptionally, consistent with previous reports (*17, 32*). Interestingly, we also found that genes with poor protein-mRNA correlations (median *ρ* of −0.2 to 0.2 in 200-gene windows) were enriched in GO-slim terms related to CCM (“carbohydrate metabolic process”, “cellular respiration”, and “generation of precursor metabolites and energy”; Fig 6B), indicating a distinct role of protein translation in regulating CCM gene expression which reflects the unique and complex mechanisms that regulate this important part of metabolism (*8*).

**Fig 6.**
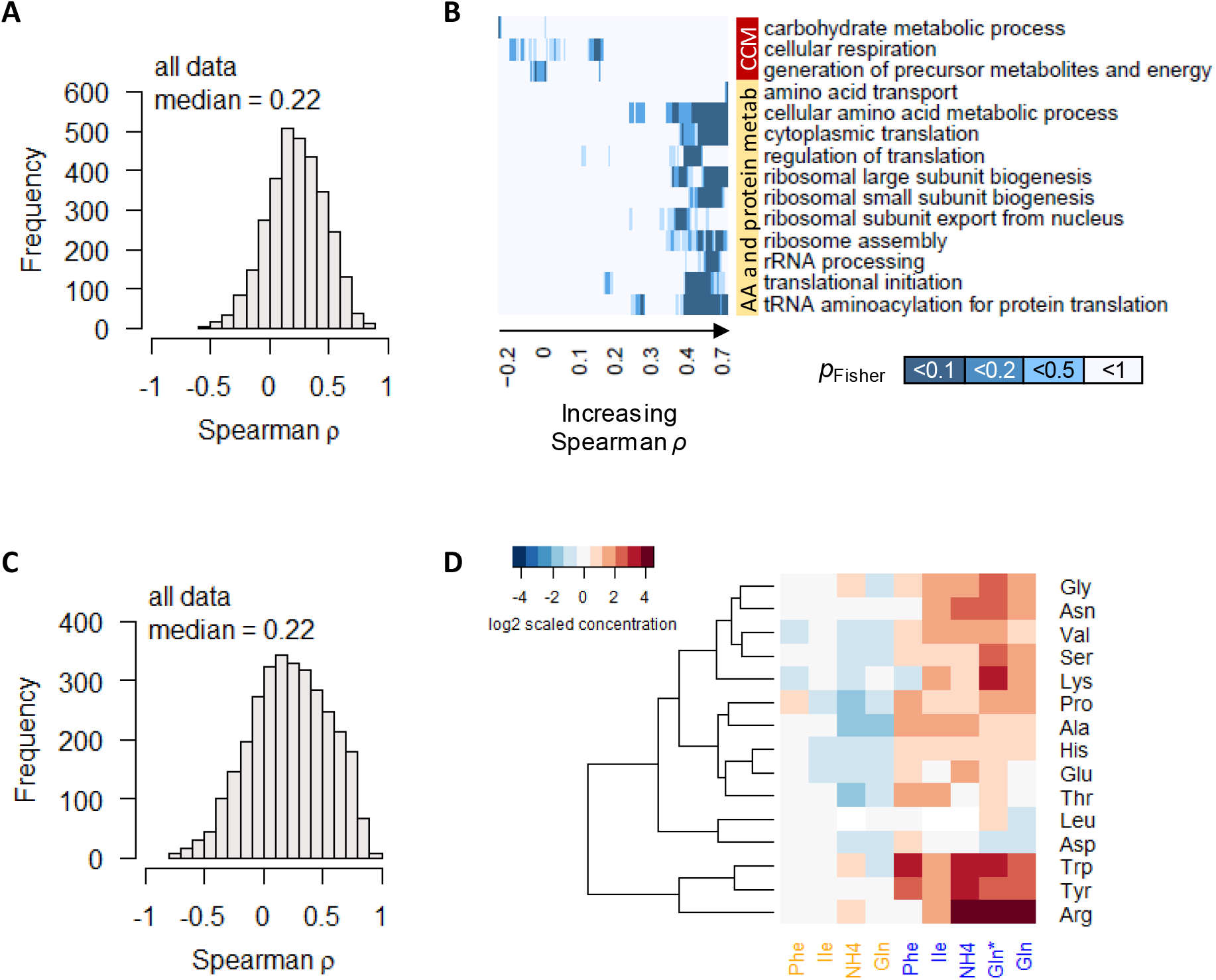
Global control of protein synthesis. A. Spearman correlation of absolute-quantitative mRNA and protein abundances for each gene is calculated using data from both the GR and the AA experiments combined, and the distribution is shown demonstrating overall poor correlation. B. Enrichment of GO-slim terms in 200-gene sliding windows of increasing Spearman correlations was analyzed by two-tailed Fisher’s exact test. Shown are GO-slim terms related to central carbon metabolism (CCM), and amino acid (AA) and protein metabolism, with at least one sliding-window with *p*_Fisher_<0.05, indicating that genes with poor protein-mRNA correlations are enriched in CCM, while genes with good protein-mRNA correlations are enriched in AA and protein metabolism. C. Spearman correlation of absolute-quantitative mRNA and protein abundances for each gene is calculated using data from both the GR and the AA experiments combined, and the distribution is shown demonstrating overall poor correlation. D. Relative intracellular concentrations of amino acids in chemostat cultures constituting the AA experiments. Orange colored column labels (first 4 columns) indicate nitrogen-limited cultures; blue colored column labels (last 5 columns) indicate carbon-limited cultures. Column labels indicate the nitrogen source. In C-limited conditions (blue), the “Gln” condition and the “Gln*” condition differ in the concentration of Gln and glucose in the chemostat feed media; see Table S1 for full details. Here 15 amino acids are shown; Table S4 contains also the measurements of Gln, Ile, and Phe, which are not plotted here as the concentrations of these 3 amino acids, in the cultures where they are the non-limiting nitrogen source, are very high and skews the colour scale of the heatmap.

### Relative protein abundance is not controlled by relative mRNA abundance

One question that arises from the AA experiments is why 96% of transcripts consistently exhibited large differential expression (Fig 3B), which for most genes do not translate into changes in protein abundance in a concordant way (Fig 5D). One possibility is that relative mRNA expression provides information on how to partition ribosomes in the cell to translate different proteins, while the absolute abundance of mRNA may be irrelevant. Numerous ribosome profiling studies (*33, 34*) have shown that ribo-seq data are highly correlated with RNAseq data, suggesting that ribosomes are binding to mRNA according to relative mRNA quantities. In the AA experiments, ribosomal proteins constituted a constant ~27% (mol/mol) of the proteome (Fig 5F), which seems to suggest that ribosome availability may place an overall constraint on protein synthesis and hence protein abundance. However, the correlation of relative-quantitative mRNA abundance (% mol/mol of total mRNA) with relative-quantitative protein abundance were also poor (median *ρ* of 0.22; Fig 6C). Thus we conclude that protein abundance was not simply controlled by relative mRNA abundance and constrained by total ribosome content. Rather, transcription and translation appeared to be two distinct points of control that were independently modulated via metabolic cues, which combined to produce proteins at the observed abundances.

To search for such metabolic cues, we measured the intracellular concentrations of 18 amino acids in the AA experiments, and found that most amino acids were accumulated by several fold when cells were grown under carbon-limitation on preferred nitrogen sources (Fig 6D and Table S4). Our data therefore suggest a model of global gene expression control by metabolic parameters, whereby protein abundance was tuned primarily through gene-specific translational regulation. When the growth condition improved such that excess amino acids could accumulate (Fig 6D), the increase in Gly and Ser promoted purine synthesis (Fig 4A) which allowed for an increase in transcript abundance for most genes (Fig 3B and Fig 4C), leading to the observed uncoupling of transcript abundance and protein abundance (Fig 5D). Of note, CCM genes (as well as amino acid metabolism genes) were enriched in the 4% of genes that did not show this characteristic change in transcript abundance (Fig 3C-D), suggesting that the transcript abundance of these genes were actively controlled by the cell, again highlighting that CCM gene expression is regulated by mechanisms that are distinct from global physiological control. Together, our overall model of global gene expression regulation by growth rate and metabolic parameters is presented in Fig 7.

**Fig 7.**
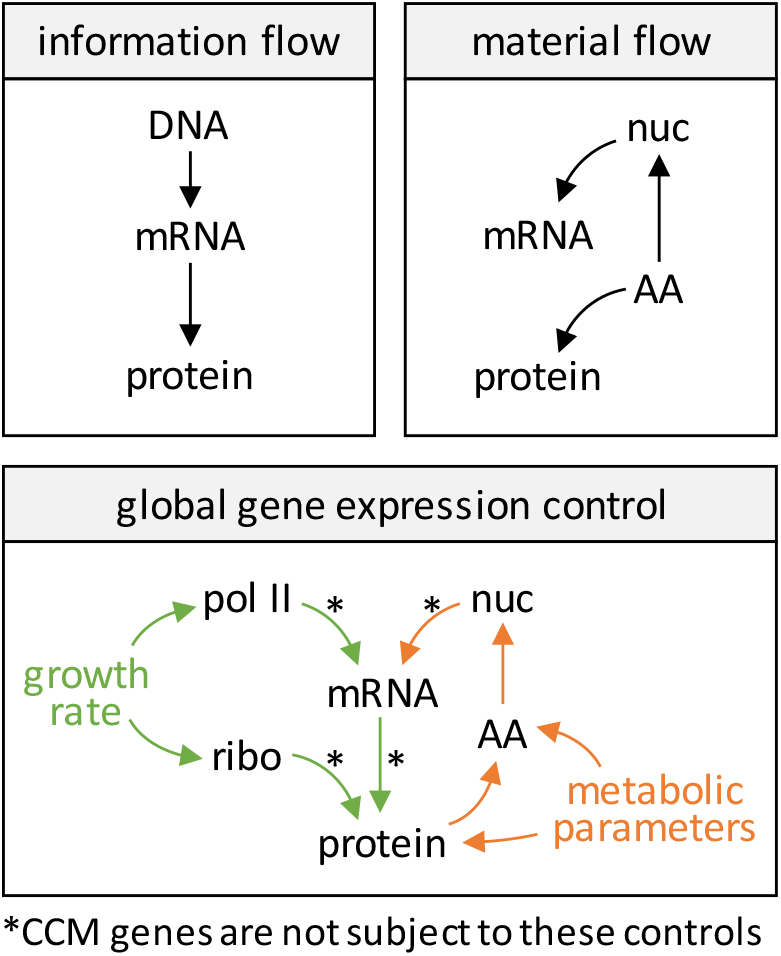
Model of information flow, material flow, and global control of material abundance in the central dogma. Nuc, nucleotides; AA, amino acids; pol II, RNA polymerase II; ribo, ribosome; CCM, central carbon metabolism. In material flow, intracellular amino acids satisfy both protein synthesis as well as nucleotide and mRNA synthesis. In global gene expression control, intracellular amino acid begins to accumulate when protein synthesis is satisfied, thus protein abundance controls amino acid abundance. Increased amino acid abundance then leads to increased nucleotide and mRNA synthesis, hence high amino acid abundance is associated with lower protein translation rate (*k_sP_*).

### Augmenting gene expression profiling analyses

With an understanding of the global effects of the cell growth rate and amino acid metabolic parameters on gene expression (Fig 7), these effects can be factored into gene expression analyses to allow the extraction of gene expression programs related specifically to the perturbation of interest (Fig 1A). Here we showcase two such applications of our findings, first using a dataset in a distantly related yeast, *S. pombe,* and second using a dataset from a human cancer cell model.

Marguerat *et al* (*19*) have shown that in *S. pombe,* gene expression in quiescent cells was characterized by a drastic downregulation of the total transcriptome compared to proliferating cells, with >85.3% transcripts being lowered by >2-fold. However, there is an inherent difference in growth rate between proliferating and quiescent cells, and quiescence was induced experimentally by nitrogen starvation (*19*). Together, we expect these differences in growth rate and metabolic parameters to influence the abundance of >90% of transcripts. To account for genes whose expression was independent of regulation by growth rate or nitrogen limitation, we took a coarsegrained approach (*9*) and considered all genes involved in CCM (PomBase GO slim term “carbohydrate metabolic process”) (*35*) to be expressed at a constant level. We found that for most transcripts, the difference in abundance between proliferating and quiescent cells were largely in line with the difference in growth rate (Fig 8A). After accounting for the global effect of growth rate on transcript expression, quiescence *per se* induced a specific gene expression program by upregulating 707 transcripts (14%) and downregulating 574 transcripts (11%). In contrast, protein translation dynamics was extensively remodeled by quiescence, with 90% of *k_sP_* being upregulated (Fig 8B). These results suggest that protein translation dynamics should be emphasized in future studies of quiescence in *S. pombe,* more so than transcriptome profiling.

**Fig 8.**
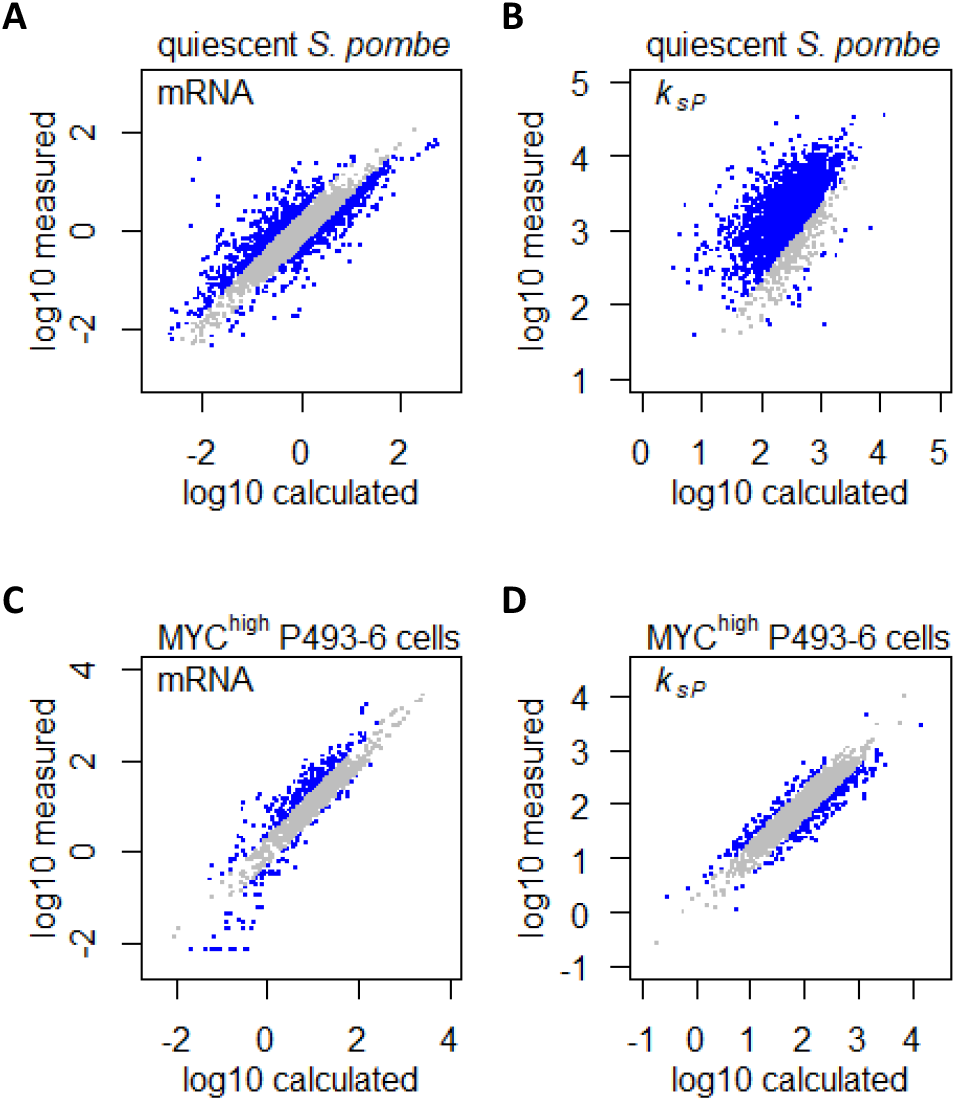
Augmenting gene expression profiling analyses. A. Measured mRNA abundances in quiescent *S. pombe* are compared to the calculated mRNA abundance, given the mRNA abundance in proliferating *S. pombe* and the difference in cell growth rate and metabolic parameters. Grey data points are those where calculated and measured values agree within 2-fold change (−1 < log_2_ < 1). B. Protein translation rate *k_xP_* calculated in quiescent *S. pombe* cells are compared to the protein translation rate *k_sP_* calculated based on protein and mRNA abundance in proliferating *S. pombe* and the difference in cell growth rate and metabolic parameters. Grey data points are those where calculated and measured values agree within 2-fold change (−1 < log_2_ < 1). C. Measured mRNA abundance in MYC-overexpressing (MYC^high^) P493-6 cells are compared to the calculated mRNA abundance, given the mRNA abundance in P493-6 cells without MYC overexpression (MYC^low^) and the difference in cell growth rate and metabolic parameters. Grey data points are those where calculated and measured values agree within 2-fold change (−1 < log_2_ < 1). D. Protein translation rate *k_XP_* calculated in MYC-overexpressing (MYC^high^) P493-6 cells are compared to the protein translation rate *k_sP_* calculated based on protein and mRNA abundance in P493-6 cells without MYC overexpression (MYC^low^) and the difference in cell growth rate and metabolic parameters. Grey data points are those where calculated and measured values agree within 2-fold change (−1 < log_2_ < 1).

We then applied our findings in a human cancer cell model, to showcase that changes in gene expression caused by oncogenic perturbation can be uncoupled from those changes that are induced by global physiological parameters such as increased growth rate and altered metabolic parameters in cancer cells. We mined gene expression datasets in the B-cell line P493-6 (*36, 37*), which carries a conditional *c-Myc* allele that can be experimentally turned on (MYC^high^), mimicking overexpression of the MYC transcription factor which is the driving oncogenic event in Burkitt’s lymphoma (*38*). Lin *et al (36*) quantified the absolute abundance of 1,263 transcripts in the P493-6 cell system, and found that 707 transcripts (56%) were upregulated by >2-fold in MYC^high^ cells. Similarly, Feist *et al (37*) quantified the absolute abundance of 1,662 proteins, and found that 871 proteins (52%) were upregulated by >2-fold in MYC^high^ P493-6 cells. However, MYC overexpression in P493-6 also leads to faster cell growth (*38*) and dependency on Gln as a nitrogen source (*37*), both of which could influence gene expression globally. We therefore sought to uncouple the effects of these physiological factors from the specific effects of MYC overexpression, which would help to elucidate direct MYC targets. Taking a similar coarse-graining approach as before, we considered all CCM genes (AmiGO 2 term “cellular carbohydrate metabolic process”) (*39*) to be expressed independently of global physiological control of gene expression. Our analysis indicated that the majority of the upregulated transcripts could be accounted for by changes in the cell growth rate and metabolic parameters (Fig 8C), while MYC overexpression specifically upregulated 309 transcripts (24%) and downregulated 167 transcripts (13%). Similarly, protein translation dynamics in MYC^high^ P493-6 cells were also largely accounted for by changes in the cell growth rate and metabolic parameters (Fig 8D), with MYC overexpression causing the up-and down-regulation of *k_sP_* for 101 (6%) and 212 (13%) proteins, respectively. Finally, we observed that genes with upregulated mRNA abundance generally had lowered *k_sP_*, and vice versa (Supplemental Fig 6), indicating that protein translation plays a buffering role in MYC-overexpressing cancer cells, producing dampened effects on protein abundance compared to changes in transcript abundance, which will be an important consideration in efforts to develop therapeutics based on transcriptome profiling studies.

## Discussion

Gene expression regulation is necessarily coupled to the physiological state of the cell, which controls global cellular parameters such as the abundance of RNA polymerases and ribosomes (*4*), as well as the sizes of transcriptomic and proteomic reserves (*8, 40*). Here we quantified the global effects of cell growth rate and amino acid metabolic parameters on gene expression, by analyzing an orthogonal multi-omics dataset consisting of paired absolute-quantitative transcript-protein abundances of 22 stead-state yeast cultures. Overall >90% of genes are regulated by the growth rate and/or amino acid metabolism of the cell at both the transcript and protein levels, suggesting widespread confounding effects in gene expression profiling studies where these factors are commonly neglected. The availabilities of RNA polymerase II and ribosomes are major contributors of differential gene expression at different cell growth rates, while the availabilities of amino acids and nucleotides underly the global gene expression changes associated with different metabolic parameters (Fig 7). Additionally, transcription and translation are regulated in a coordinated manner in response to growth rate, but are regulated in distinct ways in response to metabolic parameters, highlighting the complex interaction between different physiological parameters in the global regulation of gene expression.

Previously we have shown that the cell growth rate determines the allocation of the proteome and transcriptome to different processes, including the relative abundance of RNA polymerases and ribosomes (*8, 21*). Our results here indicate that these growth rate-dependent changes in the transcription and translation machineries directly influences the absolute abundance ~90% of genes. However, the expression of genes involved in CCM are largely independent of the cell growth rate. Of note, this means that the allocation of the transcriptome and proteome (i.e. the relative abundance normalized to total transcripts and protein abundance, respectively) to CCM would decrease with growth rate (*41, 42*), highlighting this important distinction between absolute and relative quantification of gene expression. Indeed, in their seminal paper on this subject, Brauer *et al.* (*11*) have shown using relative-quantitative data that the transcript expression of 25% of genes were linearly correlated with growth rate, which included both upregulation and down regulation. Further, genes that were downregulated with growth rate were enriched in carbon metabolic processes and peroxosomal functions (*11*). Here we showed that, in absolute-quantitative terms, CCM genes were expressed at constant levels independent of the cell growth rate. However, with 90% of genes having a positive relationship with the cell growth rate, in relative-quantitative terms CCM genes would appear to be negatively regulated by the cell growth rate, consistent with the previous report (*11*).

In addition to the cell growth rate, we showed that the amino acid metabolic parameters of the cell is also a global regulator of gene expression, consistently modulating the overall abundance of 96% of transcripts. We found that gene expression in response to amino acid metabolic parameters is not dependent on how much RNA polymerases and ribosomes are present in the cell, but instead is closely associated with the availability of amino acids and nucleotides. Genes demonstrating the strongest uncoupling of transcription and translation as two distinct points of regulation-i.e. genes whose transcript and protein abundances were poorly correlated – were particularly enriched for CCM processes, consistent with previous indications that this central metabolic pathway is regulated by complex mechanisms to ensure robust output (*8*). We further showed that protein abundance is unlikely to be controlled by relative mRNA abundance, in line with recent large-scale proteomics studies showing that the organizing principle of the cell proteome can be substantially different from that of the transcriptome (*43*). Our data therefore suggests a model of gene expression regulation by metabolic parameters, whereby protein abundance is tuned primarily through gene-specific translational regulation; meanwhile, transcript abundance increases (or decreases) with the availability of amino acids and nucleotides, with the exception of a small number of genes enriched in CCM and amino acid metabolism (Fig 7).

In many experimental systems (e.g. drug treatment, wildtype vs mutant, expression of transgenes), the physiological state of the cell is different between experimental conditions but is difficult to control for. Recent approaches to gain such control include the development of isogrowth gene expression profiling, whereby cell cultures treated with a 2D-drug gradient are sampled along a line of constant growth (the isobole), in order to dissect the gene expression changes without the confounding factor of changing growth rate (*44*). This methodology is however technically challenging and difficult to scale up. We show here that effects of growth rate and metabolism can be disentangled with the results presented in this study. Of note, it is particularly important to take these effects into consideration when the fold-change in gene expression is small (i.e. a few fold), as very large changes (e.g. by an order of magnitude or more) are unlikely to be regulated by physiological parameters of the cell (*4*). One application of this is the gene expression changes in human cancer cells induced by overexpression of the oncogene MYC, which has been suggested to play an amplifier role to drive small increases in all expressed transcripts (*36*), but this effect has been debated (*45*). Our analysis indicates that the increase in gene expression is in line with altered cell physiology during MYC-driven oncogenic transformation (*37, 38*), which then allowed MYC-specific effects of transcript expression and protein translation to be identified. Thus, by understanding the global effects of gene expression exerted by the physiological state of the cell, our results provide a framework to analyze gene expression profiles to gain novel biological insights, a key conceptual goal of systems biology.

## Materials and methods

### Culture conditions

The yeast *Saccharomyces cerevisiae* CEN.PK113-7D (MATa, MAL2-8c, *SUC2*) was used for all experiments. Cells were stored in aliquoted glycerol stocks at −80 °C. Chemostat experiments were carried out in DASGIP 1L bioreactors (Jülich, Germany) equipped with off-gas analysis, pH, temperature and dissolved oxygen sensors. Chemostat experiments were carried out at 30°C, pH 5, working volume 0.5 L, aeration 1 vvm, pO_2_ >30%, agitation speed 800 rpm. The glucose and nitrogen source concentrations are in Table S1. Additional media components are: KH_2_PO_4_, 3 g L^-1^; MgSO_4_·7H_2_O, 0.5 g L^-1^; trace metals solution, 1 ml L^-1^; vitamin solution, 1 ml L^-1^; antifoam, 0.1 ml L^-1^. The trace metals solution contained: EDTA (sodium salt), 15.0 g L^-1^; ZnSO_4_·H_2_O, 4.5 g L^-1^; MnCl_2_·2H_2_O, 0.84 g L^-1^; CoCl_2_·6H_2_O, 0.3 g L^-1^; CuSO_4_·5H_2_O, 0.3 g L^-1^; Na_2_MoO_4_·2H_2_O, 0.4 g L^-1^; CaCl_2_·2H_2_O, 4.5 g L^-1^; FeSO_4_·7H_2_O, 3.0 g L^-1^; H_3_BO_3_, 1.0 g L^-1^; and KI, 0.10 g L^-1^. The vitamin solution contained: biotin, 0.05 g L^-1^; p-amino benzoic acid, 0.2 g L^-1^; nicotinic acid, 1 g L^-1^; Ca-pantothenate, 1 g L^-1^; pyridoxine-HCl, 1 g L^-1^; thiamine-HCl, 1 g L^-1^ and myo-inositol, 25 g L^-1^.

### Sampling from bioreactor

The dead volume was collected with a syringe and discarded. For transcriptome sampling, biomass was collected from the reactor with a syringe and injected into chilled 50 ml Falcon tubes filled with 35 mL crushed ice. Samples were centrifuged for 4 min at 3,000 x g at 4 °C; cell pellets were washed once with 1 mL of chilled water, transferred into Eppendorf tubes, flash frozen in liquid nitrogen, and stored at −80 °C until analysis. For proteome sampling, biomass was collected from the reactor with a syringe and injected into 50 ml Falcon tubes chilled on ice. Samples were centrifuged for 4 min at 3,000 x g at 4 °C; cell pellets were washed once with 20 ml of chilled dH_2_O, washed again with 1 ml of chilled water, transferred into Eppendorf tubes, flash frozen in liquid nitrogen, and stored at −80 °C until analysis. For intracellular amino acid analysis, biomass was collected from the reactor with a syringe and injected into −80 °C MetOH at 10-times the volume of the sample. Sample were centrifuged to 4 min at 3,000 x g at −20 °C, pellets were flash frozen in liquid nitrogen and stored at −80 °C until analysis. Biomass determination was done by filtration of the culture broth on preweighed filter paper, drying in a microwave at 360 W for 20 min, and desiccating in a desiccator for >3 days.

### RNA sequencing

RNA was extracted using Qiagen RNeasy Mini Kit (Qiagen, Hilden, Germany) according to manufacturer’s protocol. RNA integrity was examined using a 2100 Bioanalyzer (Agilent Technologies, Santa Clara, CA). RNA concentration was determined using a Qubit RNA HS Assay Kit (Thermo Fisher, Waltham, MA). The Illumina TruSeq Stranded mRNA Library Prep Kit (Illumina, San Diego, CA) was used to prepare mRNA samples for sequencing. Paired-end sequencing (MID Output 2×75 bp) was performed on an Illumina NextSeq 500 (Illumina, San Diego, CA). Reads were quality controlled, mapped to the *S. cerevisiae* reference genome (Ensembl R64-1-1), and counted using the nf-core RNAseq pipeline (SciLifeLab, Stockholm, Sweden), available at https://nf-co.re/rnaseq.

### Quantitative proteome measurements

LC-MS based proteomic analysis was performed largely as described in our previous publication (*8*). The Supplementary Methods section contains the detailed description of the protocol. Briefly, cell pellets were lysed via bead beating in presence of 2% sodium dodecyl sulfate (SDS) and an aliquot from each sample was digested using Pierce MS grade trypsin (Thermo Fisher), labeled with TMT 10plex isobaric tags (Thermo Fisher), the labeled and combined samples were fractionated into 20 fractions each using basic-pH reversed-phase chromatography and analyzed on an Easy-nLC 1200 chromatography system coupled to an Orbitrap Fusion Tribrid mass spectrometer (both Thermo Fisher) for the relative protein quantification. The pooled reference mixture that consisted of an equal amount of protein from each sample was spiked with UPS2 Proteomics Dynamic Range Standard (Sigma-Aldrich, Saint-Louis, MO) and digested with trypsin, followed by fractionation into 10 fractions and LC-MS analysis for the estimation of the absolute protein amounts via IBAQ (*18*).The LC-MS files were processed in Proteome Discoverer 2.2 (Thermo Fisher), including the database search with Mascot 2.5.1 (Matrix Science, London, United Kingdom) against the *S. cerevisiae* ATCC 204508 / S288c) reference proteome from Uniprot (February 2018, 6049 sequences) and TMT reporter ion quantification. For the IBAQ label-free data, the *S. cerevisiae* database was supplemented with the sequences of the 48 UPS2 proteins and the Minora Feature Detector in Proteome Discoverer 2.2 was used for the precursor ion quantification; the resulting protein abundances were exported and the linear regression was calculated between the known log10 UPS2 concentrations and the log10 IBAQ abundances for the corresponding proteins. The resulting regression coefficients were used to estimate the absolute concentrations of the yeast proteins.

### Intracellular amino acid measurements

To extract intracellular amino acid, 1-2 ml of boiling 75% ethanol was poured directly onto the cell pellets. Samples were vortexed for 1 min, boiled for 3 min, and placed on ice until cool. Samples were centrifuged for 15 min at 13,000 x g at 4 °C. Supernatant containing intracellular amino acids were collected and stored at −20 °C until labeling and analysis. Amino acids were labeled using the SCIEX aTRAQ Reagents Application Kit (Danaher, Washington DC) with aTRAQ Reagent Δ8, and mixed with internal standards pre-labeled with aTRAQ Reagent Δ0, according to manufacturer’s protocol. Samples were analyzed using a SCIEX QTRAP 6500+ system (Danaher, Washington DC) with a Nexera UHPLC system (Shimadzu, Japan), on a SCIEX AAA column (150 x 4.6 mm) at an oven temperature of 50 °C. Mobile phase A contained 0.1% formic acid and 0.01% heptafluorobutyric acid in water; mobile phase B contained 0.1% formic acid and 0.01% heptafluorobutyric acid in methanol. The flow rate was 0.8 mL min^-1^, with a gradient profile was as follows: 0 min, 2% B; 6 min, 40% B; 10 min, 40% B; 11 min, 90% B; 12 min, 90% B; 13 min, 2% B; 18 min, 2% B. The retention time, precursor (Q1 m/z) and product (Q3 m/z) for each amino acid and internal standard were as described by the manufacturer’s protocol. The mass spectrometer was set to monitor the transitions with the following ion source parameters: CUR (curtain gas) 30, CAD (collision activated dissociation) MED, IS (ionization voltage) 5500, TEM (temperature) 500, GS1 (source gas 1) 60 and GS2 (source gas 2) 50; and with compound parameters: DP (declustering potential) 30, EP (entrance potential) 10, CE (collision energy) 30 and CXP (collision cell exit potential) 5.

### Data processing and analysis

For transcriptomics, the absolute concentrations of 31 transcripts with >10 FPKM, and covering the entire dynamic expression range, were measured using lysates of *S. cerevisiae* CEN.PK 113-7D cells from the J. Nielsen lab (Chalmers University of Technology, Sweden). Linear regression between the absolute concentrations of these mRNAs and their corresponding FPKM values from RNAseq were performed to obtain the slope and y-intercept, which were used to quantify all mRNA in this study. The calculated mRNA abundance was then scaled to the total RNA content measured by Qubit RNA HS Assay Kit (Thermo Fisher, Waltham, MA). For proteomics, detailed data processing parameters are described in the Supplementary Methods.

Protein translation rate *k_sP_* (*17,18*) for each protein *j* was calculated by:

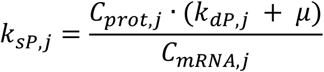

The gene-specific protein degradation rate (*k_dP_*) for *S. cerevisiae* was mined from Lahtvee *et al* (*17*). Protein degradation rate for *S. pombe* was mined from Christiano *et al* (*46*). Protein degradation rate for a cancer cell line (HeLa) was mined from Boisvert et al (*47*). Where the *k_dP,j_* was unavailable in the mined datasets, the median of all *k_dP_* in the dataset was used.

### Data and code availability

Processed quantitative transcriptomics and proteomics data are in Supplementary Table 2 and 3. Processed intracellular amino acid concentrations are in Supplementary Table 4. Raw RNAseq data are available at ArrayExpress, accession E-MTAB-9117 (*48*). The mass spectrometry proteomics data are deposited to the Proteome Xchange Consortium via the PRIDE (*49*) partner repository with dataset identifier PXD021218 (*50*). GO-slim terms for *S. cerevisiae* genes are available from the Saccharomyces Genome Database (*25, 51*). GO-slim terms for *S. pombe* genes are available from PomBase (*35, 51*). GO terms for human genes are available from AmiGO 2 (*39, 51*). All other supporting data and custom code are available on request from the corresponding author.

## Supporting information

reviewer login for data depositories

supoplemental tables

## Acknowledgements

We thank the National Bioinformatics Infrastructure Sweden (NBIS) for discussions. We thank Pannipa Pornpitakpong and Alexandra Hoffmeyer (Technical University of Denmark) for conducting RNA sequencing. The Proteomics Core Facility of the University of Gothenburg is grateful to the Inga-Britt and Arne Lundbergs Forskningsstiftlese for the donation of the Orbitrap Fusion Tribrid MS instrument. This research was supported by funding from the Novo Nordisk Foundation (grant number NNF10CC1016517) and the Knut and Alice Wallenberg Foundation.

