## Supplementary material for "Quantifying absolute gene expression profiles reveals distinct regulation of central carbon metabolism genes in yeast": reviewer login for data depositories

### Transcriptomics data depository – reviewer access

Access at: <http://www.ebi.ac.uk/arrayexpress/experiments/E-MTAB-9117>

Project accession: E-MTAB-9117

Username: Reviewer_E-MTAB-9117

Password: vpxsVL26

### Proteomics data depository – reviewer access

Access at: <https://www.ebi.ac.uk/pride/archive/login>

Project accession: PXD021218

Username:

Password: x8KSbPxh
