## supplemental figures and methods for "Quantifying absolute gene expression profiles reveals distinct regulation of central carbon metabolism genes in yeast"

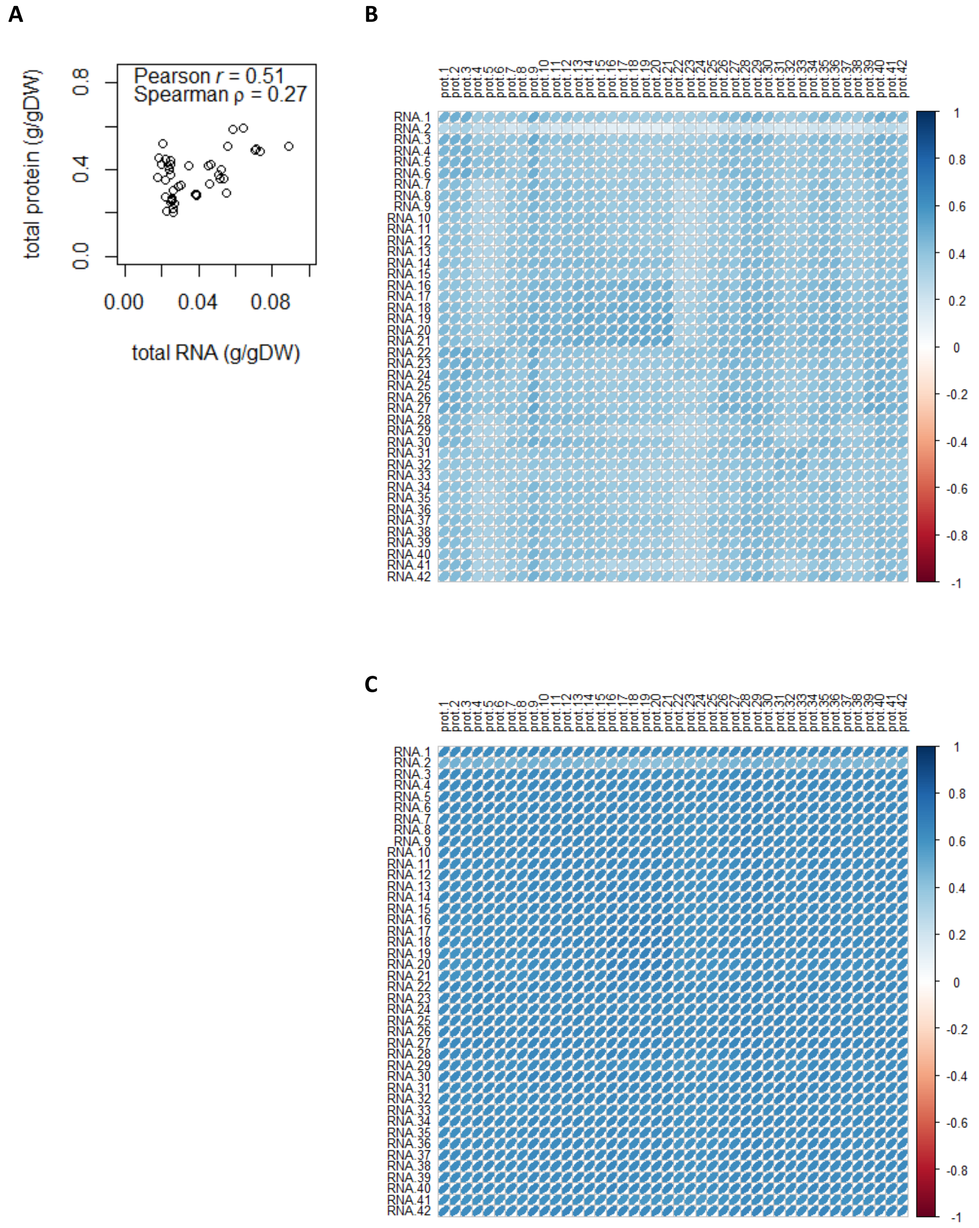


**Supplemental Fig 1. Total and sample-wise protein-mRNA correlations.**

A. Correlation between total protein content and total mRNA content (g/gDW).

B. Sample-wise proteome-transcriptome Pearson correlations. The median Pearson *r* is 0.40. See Table S1 for sample ID and full details of chemostat conditions.

C. Sample-wise proteome-transcriptome Spearman correlations. The median Spearman *ρ* is 0.63. See Table S1 for sample ID and full details of chemostat conditions.

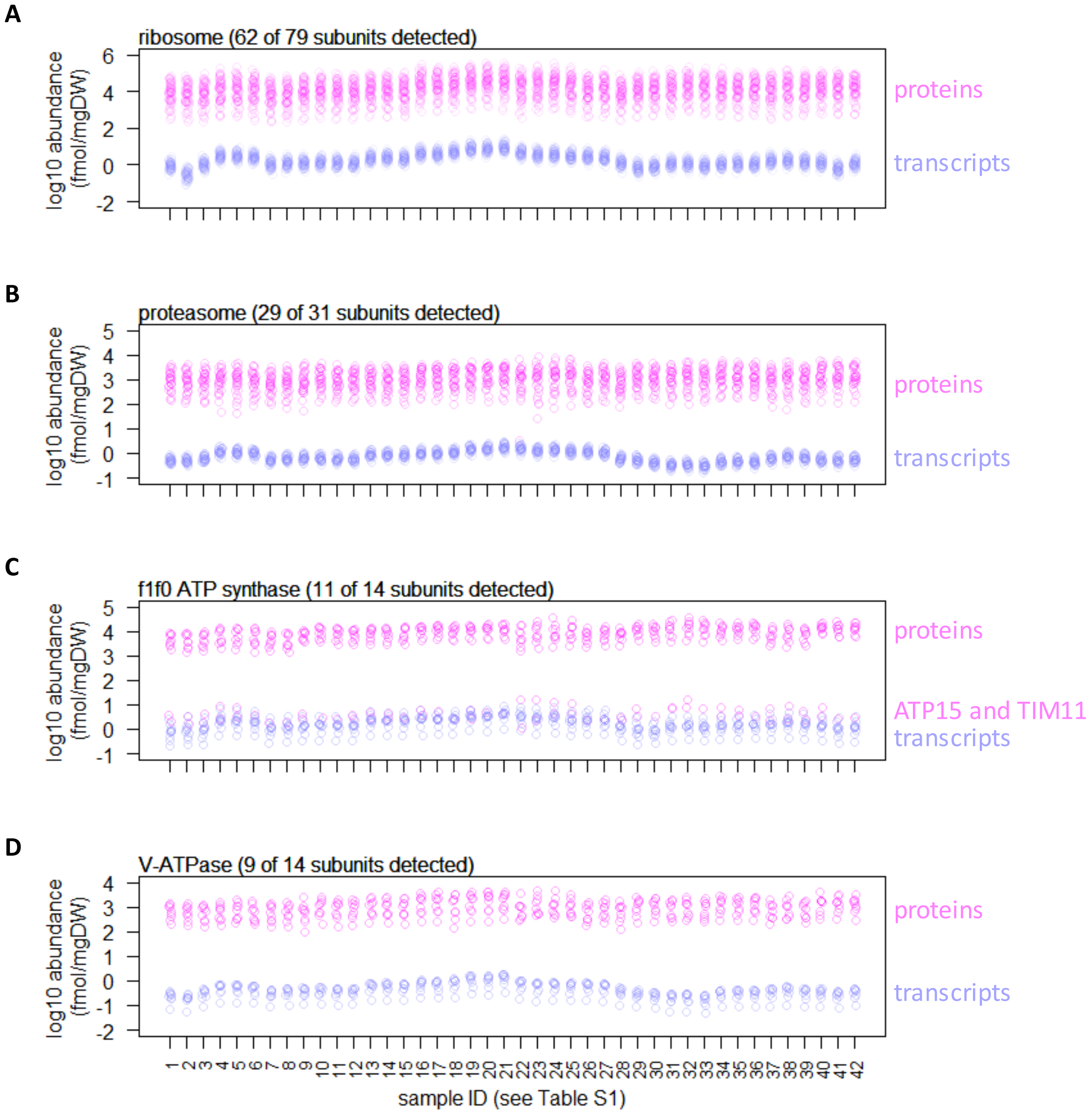


**Supplemental Fig 2. Measured absolute quantity of proteins (pink) and transcripts (blue), for subunits of protein complexes as indicated.**

For ribosomes (panel A), the abundance of pairs of paralogs (e.g. RPS14A and RPS14B) are summed.

For f1f0 ATP synthase (panel C) and V-ATPase (panel D), shown are absolute abundance of the subunits after normalization to the stoichiometry (e.g., in f1f0 ATP synthase, ATP1 and ATP2 are resent at 3 subunits per complex; the absolute abundance is therefore divided by 3 before plotting in the graph above).

For f1f0 ATP synthase (panel C), the protein abundance of ATP15 and TIM11 are ~4 orders of magnitude lower than the abundance of other subunits, likely reflecting the difficulty in extracting and quantifying membrane-embedded proteins.

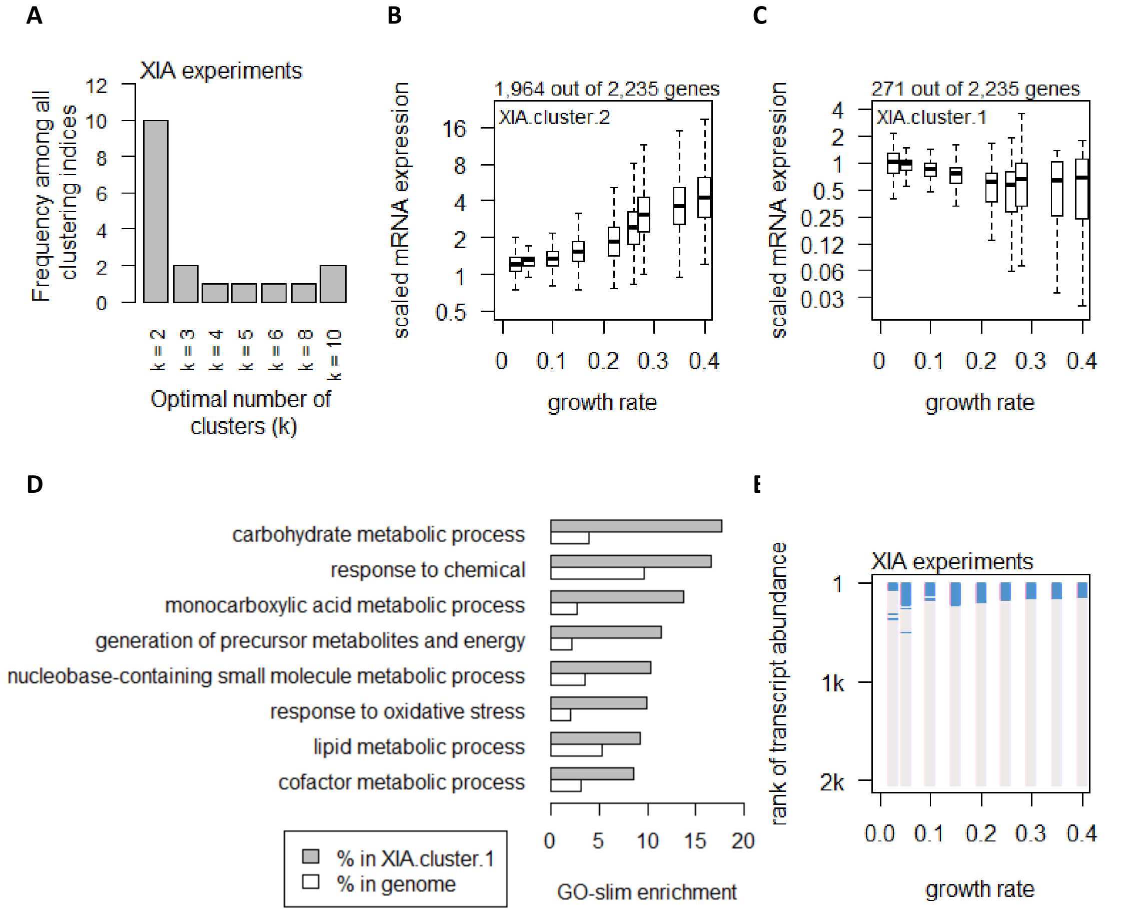


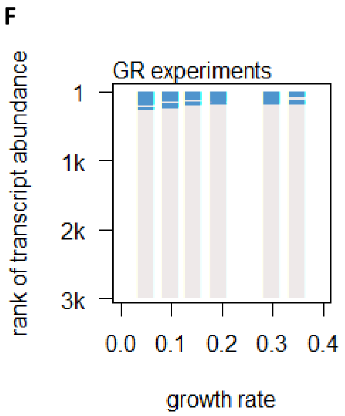


**Supplemental Fig 3. Global control of transcript abundance by growth rate.**

A. Using 25 clustering indices, we found that most indices suggest an optimal number of 2 clusters for transcript abundance in the XIA experiments.

B. Abundance of transcripts in XIA.cluster.2. Center line, median; box limits, upper and lower quartiles; whiskers, 1.5x interquartile range.

C. Abundance of transcripts in XIA.cluster.1. Details are as panel B.

D. GO-slim enrichment of genes in XIA.cluster.1 showing enrichment in GO-slim terms related to CCM.

E. Enrichment of CCM genes in 200-gene sliding windows of increasing transcript abundance in the XIA experiments is shown by two-tailed Fisher's exact test. Blue color indicates $p_{\text{Fisher}}$<0.05.

F. Enrichment of CCM genes in 200-gene sliding windows of increasing transcript abundance in the GR experiments. Details are as panel E.

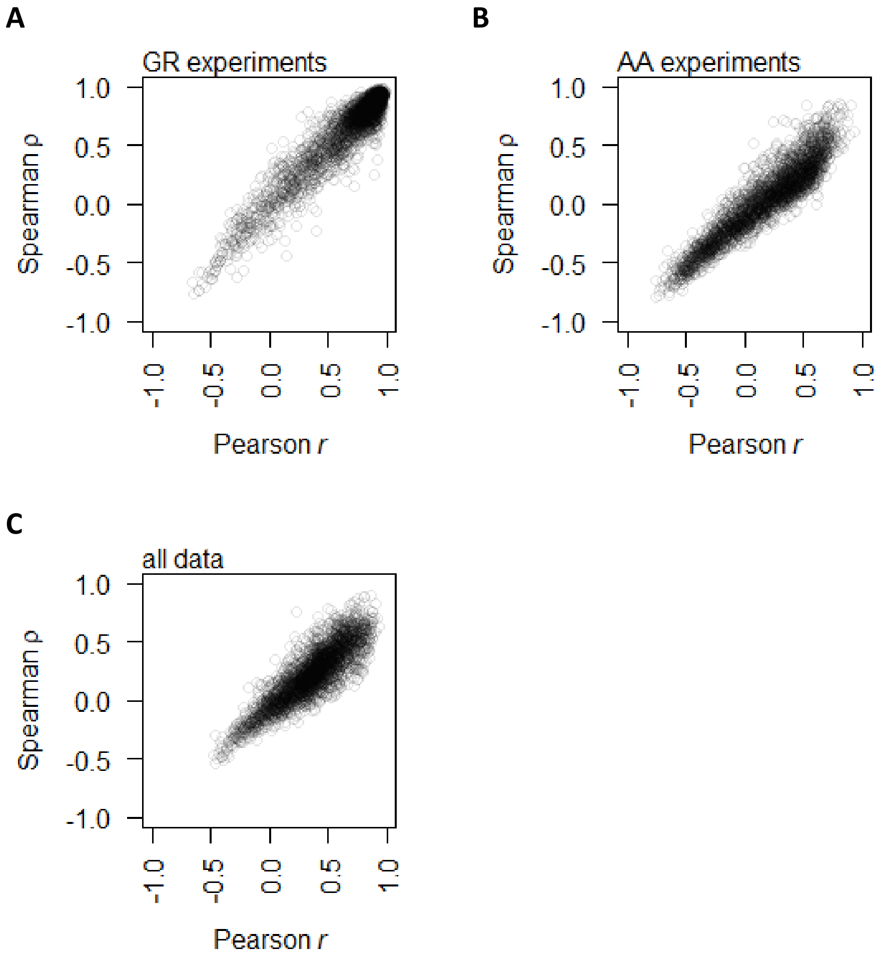


**Supplemental Fig 4. Correlation between protein and mRNA abundance for each gene showing that Spearman and Pearson correlations track well with each other. This study uses Spearman correlations for analyses since the underlying data distribution is not always normal, however previous literature used Pearson correlations.**

A. Spearman $\rho$ and Pearson $r$ between protein and mRNA abundance for each gene in the GR experiments alone.

B. Spearman $\rho$ and Pearson $r$ between protein and mRNA abundance for each gene in the AA experiments alone.

C. Spearman $\rho$ and Pearson $r$ between protein and mRNA abundance for each gene with data from the two experiments combined.

A

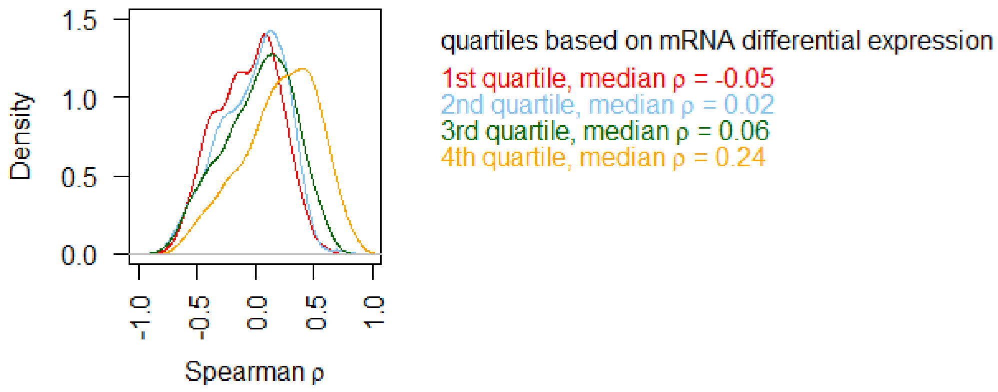


B

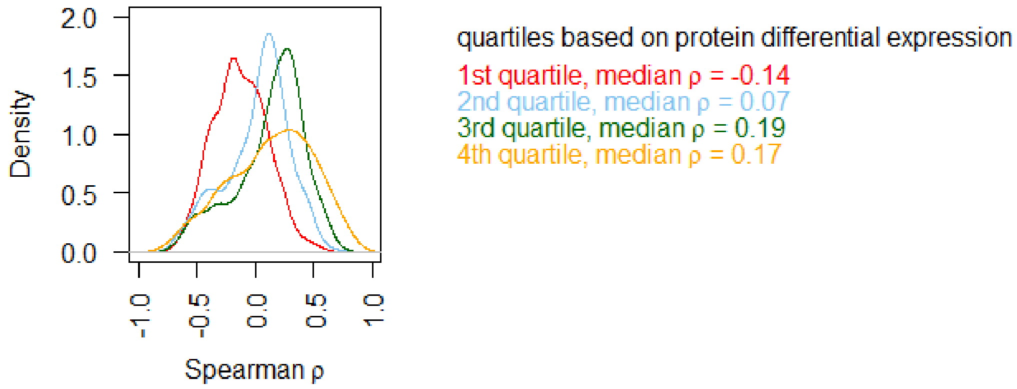


**Supplemental Fig 5. Comparing protein-mRNA correlations in the AA experiments with the range of differential expression.**

A. Genes are split into 4 groups based on quartiles of mRNA differential expression (fold-change of max to min value), and the distribution of Spearman correlation in each quartile is plotted separately.

B. Genes are split into 4 groups based on quartiles of protein differential expression (fold-change of max to min value), and the distribution of Spearman correlation in each quartile is plotted separately.

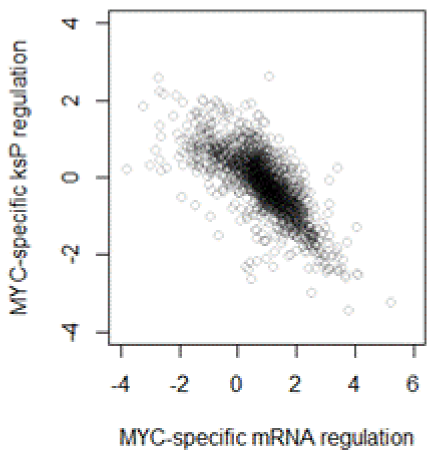


**Supplemental Fig 6. Comparison between MYC-specific regulation of mRNA abundance and protein translation efficiency ($k_{sP}$).**

### Supplemental Methods

#### Sample Preparation for Proteomic Analysis

Yeast cell pellets were suspended in the lysis buffer (2% sodium dodecyl sulfate (SDS), 50mM triethylammonium bicarbonate (TEAB)) and homogenized using a FastPrep®-24 instrument (Matrix C, 1 mm silica spheres; MP Biomedicals, OH, USA) for 5 repeated 40 seconds cycles at 6.5 m/s, with 30-60 seconds breaks in-between. The lysate was centrifuged at 21 100 xg for 20 min, the supernatant was transferred to new tubes, diluted 5 times with the lysis buffer and protein concentration was determined using Pierce™ BCA Protein Assay Kit (Thermo Fischer Scientific, Waltham, MA, USA) and the Benchmark™ Plus microplate reader (Bio-Rad Laboratories, Hercules, CA, USA) with bovine serum albumin (BSA) solutions as standards. Pooled reference sample ("Reference") has been prepared by mixing equal amounts (according to BCA measurement) from each supernatant.

For the TMT-based relative quantification, aliquots containing 25 µg of protein were taken from each experimental sample and from the pooled Reference sample; an aliquot of 50 µg of the Reference sample was spiked with 10.6 µg of the UPS2 Proteomics Dynamic Range Standard (Sigma-Aldrich, Saint-Louis, MO) for the IBAQ [1] quantification. Each sample was reduced by the addition of 2M DL-Dithiothreitol (DTT) to a final concentration of 100 mM and incubated at 56°C for 30 min. The samples were processed according to the modified filter-aided sample preparation (FASP) method [2]. In short, reduced samples were diluted to 300 µl by addition of 8M urea, transferred onto Nanosep 30k Omega filters (Pall Corporation, Port Washington, NY, USA) and washed 2 times with 200 µl of 8M urea. The free cysteine side chains were alkylated with 10 mM methyl methanethiosulfonate (MMTS) solution in digestion buffer (1% sodium deoxycholate (SDC), 50 mM TEAB) for 30 min at room temperature and the filters were then repeatedly washed with digestion buffer. Trypsin in digestion buffer was added (250 ng for TMT samples/500 ng for the IBAQ sample) and the sample was incubated at 37°C for 3h, then another aliquot of trypsin (250/500 ng) was added and incubated overnight. Digested peptides were collected by centrifugation at 9 500 xg for 20 min, followed by a wash with 20 µl of the digestion buffer and centrifugation at 9 500 xg for 20 min. Peptide samples for relative quantification were labelled using the 5 sets TMT 10plex™ isobaric reagents according to the manufacturer's instructions (Thermo Scientific), combined into 5 pooled TMT samples, concentrated using vacuum centrifugation and SDC was removed by acidification with 10% TFA and subsequent centrifugation at 16 000 xg for 10 min. The digested sample for IBAQ label-free quantification was acidified with 10% TFA and centrifuged at 16 000 xg for 10 min to remove the precipitated deoxycholic acid.

The TMT-sets were fractionated into 44 primary fractions by basic reversed-phase chromatography (bRP-LC) using a Dionex Ultimate 3000 UPLC system (Thermo Fischer Scientific, Waltham, MA, USA). Peptide separations were performed on a reversed-phase XBridge BEH C18 column (3.5 µm, 3.0x150 mm, Waters Corporation) and a linear gradient from 3% to 40% solvent B over 17 min followed by an increase to 100% B over 5 min. Solvent A was 10 mM aqueous ammonium formate at pH 10.0 and solvent B was 90% acetonitrile, 10% 10 mM ammonium formate at pH 10.00. The primary fractions were concatenated into final 20 fractions (1+21+41, 2+22+42, … 20+40), evaporated and reconstituted in 15 µl of 3% acetonitrile, 0.2% formic acid for nLC MS analysis.

The pooled sample for the label-free quantification was fractionated into 44 primary fractions that were concatenated into 10 final fractions (1+11+21+31+41, 2+12+22+32+42, … 10+20+30+40). The final fractions were evaporated and reconstituted in 15 µl of 3% acetonitrile, 0.2% formic acid for nLC MS analysis.

#### Liquid Chromatography-Mass Spectrometry Analysis

All samples were analyzed on an Orbitrap Fusion Tribrid mass spectrometer interfaced with Easy-nLC1200 liquid chromatography system (both Thermo Fisher Scientific). Peptides were trapped on an Acclaim Pepmap 100 C18 trap column (100 µm x 2 cm, particle size 5 µm, Thermo Fischer Scientific) and separated on an in-house packed analytical column (75 µm x 30 cm, particle size 3 µm, Reprosil-Pur C18, Dr. Maisch) using the linear gradients with 0.2% formic acid in water as a solvent A and 80% acetonitrile, 0.2% formic acid as solvent B with the flow of 300 nL/min.

Each TMT-labelled fraction was separated on a 90 min gradient from 5% to 35% B over 75 min, from 35% to 100% B over 5 min and 100% B for 10 min. Each fraction for the label-free IBAQ quantification was injected 3 times and analysed using the same elution gradient.

For the TMT-based relative quantification, MS scans were performed at 120 000 resolution, m/z range 380-1380 with the wide quadrupole isolation and AGC target 4e5; the most abundant precursors with charges 2 to 7 were selected for fragmentation over the 3s cycle time with the dynamic exclusion duration of 60 s. Precursors were isolated with a 0.7 Da window, fragmented by collision induced dissociation (CID) at 35% collision energy with a maximum injection time of 50 ms and AGC target 1e4, and the $MS^2$ spectra were detected in the ion trap followed by the synchronous isolation of the 10 most abundant $MS^2$ fragment ions within m/z range of 400-1400, and fragmentation by higher-energy collision dissociation (HCD) at 65%; the resulting $MS^3$ spectra were detected in the Orbitrap at 50,000 resolution with m/z range 100-500, maximum injection time 105 ms and AGC target 1e5.

For label-free quantification IBAQ experiment, MS scans were performed at 120 000 resolution, m/z range 380-1380 with the wide quadrupole isolation and AGC target 2e5; the most abundant precursors with charges 2 to 7 were selected for fragmentation over the 1s cycle time with the dynamic exclusion duration of 45 s. Precursors were isolated with a 1.2 Da window, fragmented by collision induced dissociation (CID) at 35% collision energy with a maximum injection time of 50 ms and AGC target 1e4, and the $MS^2$ spectra were detected in the ion trap.

The mass spectrometry proteomics data has been deposited to the ProteomeXchange Consortium via the PRIDE [3] partner repository with the dataset identifier PXD021218.

#### Proteomic Data Analysis

Peptide and protein identification and quantification was performed using Proteome Discoverer version 2.2 (Thermo Fisher Scientific) with Mascot 2.5.1 (Matrix Science, London, United Kingdom) as a database search engine. The baker's yeast (*Saccharomyces cerevisiae* ATCC 204508 / S288c) reference proteome database was downloaded from Uniprot (February 2018, 6049 sequences) and used for the database search on the TMT-based relative quantification files; the concatenated database containing the yeast sequences and the 48 UPS protein sequences was used for the processing of the UPS2-spiked files.

Precursor masses have been re-calibrated with the initial mass tolerance of 20 ppm prior to the main database search. For the TMT relative quantification data and the label-free IBAQ data, trypsin with 1 missed cleavage was used as a cleavage rule, MS peptide tolerance was set to 5 ppm and $MS^2$ tolerance for identification was set to 600 mmu. Variable modifications of methionine oxidation, and fixed modifications of cysteine methylthiolation were used for both sub-data sets. TMT-6 label on lysine and peptide N-termini was set as a fixed modification for the TMT-data. Percolator was used for the peptide-spectrum match (PSM) validation with the strict false discovery rate (FDR) threshold of 1%.

Precursor ion quantification was accomplished via the Minora feature detection node in Proteome Discoverer 2.2, with the maximum peak intensity values used for quantification. Abundance values of all unique and shared peptides were used for the IBAQ calculation. Abundances from the 3 technical replicates were averaged and divided by the number of theoretically observable peptides for a protein to yield the IBAQ intensity (the number of observable peptides being calculated using an in-house Python script). The known absolute amount values of the UPS2 standard proteins were used to scale the Log-transformed IBAQ intensity values versus Log-transformed protein concentration. Only UPS2 proteins with at least 2 identified unique peptides were used for scaling.

The TMT reporter ions were identified in the $MS^3$ HCD spectra with a mass tolerance of 3 milli mass units (mmu), the signal-to-noise (S/N) abundances of the reporter ions of all unique and shared peptides were used for relative quantification with the minimal average reporter S/N threshold set at  19 and the co-isolation threshold at 100. The resulting reporter abundance values for each sample were normalized within Proteome Discoverer 2.2 on the total peptide amount.
